# Dendritic synaptic integration modes under *in vivo*-like states

**DOI:** 10.1101/2024.11.18.624129

**Authors:** Cesar C. Ceballos, Rodrigo F. O. Pena

## Abstract

The neural code remains undiscovered and understanding synaptic input integration under *in vivo*-like conditions is just the initial step toward unraveling it. Synaptic signals generate fast dendritic spikes through two main modes of temporal summation: coincidence detection and integration. In coincidence detection, dendrites fire only when multiple incoming signals arrive in rapid succession, whereas integration involves summation of postsynaptic potentials over longer periods with minimal membrane leakage. This process is influenced by ionic properties, especially as the membrane potential approaches the firing threshold, where inactivating currents play a critical role. However, the modulation of temporal summation by these currents under *in vivo*-like conditions has not been thoroughly studied. In our research, we used computer simulations of a single dendritic branch to investigate how three inactivating currents — A-type potassium, T-type calcium, and transient sodium — affect temporal summation. We found that calcium and sodium currents promote integrative behavior in dendrites, while potassium currents enhance their ability to act as coincidence detectors. By adjusting the levels of these currents in dendrites, neurons can flexibly switch between integration and coincidence detection modes, providing them with a versatile mechanism for complex tasks like multiplexing. This flexibility could be key to understanding how neural circuits process information in real time.

**Significance:** Neurons consist of specialized structures that shape computation. The soma processes inputs from dendrites and communicates via its axon. Dendrites have a branched architecture. They receive excitatory signals and generate dendritic spikes. These spikes allow individual branches to convert inputs into spiking patterns prior to somatic integration. Depending on input synchrony, branches operate as coincidence detectors or integrators, which is determined by ionic currents. Our study explores how these currents affect signal summation. We found that potassium currents favor coincidence detection, while calcium and sodium currents promote integration. Cellular mechanisms finely tune these currents, enabling dendrites to adjust processing modes in response to synaptic inputs.

## Introduction

It is well known that dendrites can fire sodium, calcium and NMDA dependent action potentials (dendritic spikes) under *in vivo* conditions at higher frequency than found at the soma (Moore et al. 2017). These suggests the existence of dendritic synaptic integration and transformation into dendritic spikes as a first stage of neuronal processing (Ujfalussy et al. 2018). Understanding the neural code involves learning how dendrites process synaptic inputs and transform them into dendritic spikes via temporal summation (Stuart et al. 2015; Dembrow et al. 2022). Two key summation types, coincidence detection and integration, are critical for neural coding (Brette 2015; Ratté et al. 2013; Hong et al. 2012; Stoll et al. 2023). Coincidence detection occurs when a dendrite fires a spike after receiving multiple signals within a short interval, while integration mode allows voltage accumulation over time until a spike fires (Dembrow et al. 2015; Dembrow et al. 2022; Roome et al. 2020). Dendrites exhibit both modes, with Purkinje neurons dendrites functioning as coincidence detectors and L5 dendrites in medial prefrontal cortex acting as integrators (Dembrow et al. 2015; Roome et al. 2020). These modes influence the transmission of information via firing rates or synchronized spikes (Hong et al. 2012).

These synaptic summation modes form a continuum from coincidence detection to perfect integration (Rudolph et al. 2003; Prince et al. 2021). The specific mode of summation depends on the composition of ionic currents in the dendritic membrane, which modulate synaptic temporal properties (Azouz et al. 2000; Prince et al. 2021). Strong potassium currents are linked to coincidence detection, while sodium and calcium currents support perfect integration (Mathews et al. 2010; Farries et al. 2010; Branco et al. 2016; Prescott and De Koninck 2005). Adjusting the expression of these currents allows a range of temporal summation modes.

Dendritic inactivating currents are crucial for near-threshold synaptic summation, as shown by studies on A-type potassium currents influencing theta oscillations and temporal summation (Bourdeau et al. 2007; Rathour et al. 2016). Excitatory postsynaptic potentials (EPSPs) are amplified by T-type calcium currents, enhancing temporal precision (Fukaya et al. 2018), while transient sodium currents act at subthreshold potentials (Carter et al. 2012; Stuart et al. 2015). Further research is needed to explore inactivation’s role in dendritic properties.

A complete understanding requires examining currents with (i) established, (ii) less established, and (iii) inconsistent effects. (i) Non-inactivating currents like M-type potassium reduce EPSP amplitude and shorten decay (Hönigsperger et al. 2015; Peng et al. 2017; Liu et al. 2008), while persistent sodium currents amplify and prolong EPSPs (Ceballos et al. 2017b; Liu et al. 2008; Hsu et al. 2018). (ii) Inactivating currents, such as A-type potassium, consistently reduce EPSP amplitude (Hoffman et al. 1997; Mathews et al. 2010; Bourdeau et al. 2007; Rathour et al. 2016). (iii) Calcium currents show mixed effects: some studies report no impact on amplitude or decay (Stuart et al. 1995; González-Burgos et al. 2001; Andreasen et al. 1999; Hsu et al. 2018; Seong et al. 2014), others show effects on amplitude (Fukaya et al. 2018; Gillessen et al. 1997), only on decay (Prescott et al. 2005), or both (Liu et al. 2008; Connelly et al. 2016; Urban et al. 1998; Evans et al. 2017; Deleuze et al. 2012). Transient sodium currents also show variable effects: some studies report no effect (Connelly et al. 2016; Grunditz et al. 2008), while others show amplification of EPSPs (Carter et al. 2012; Hage et al. 2016) or Tetrodotoxin-sensitive prolonged decay (Ceballos et al. 2018). Sodium-dependent EPSP amplification has been observed *in vivo* (Hsu et al. 2018).

Most EPSP studies are not easily transferable to *in vivo* situations, where neural coding is better understood. *In vivo*, large depolarization states occur during neuronal oscillations like theta oscillations (Adam et al. 2019; Malezieux et al. 2020). These states bring the membrane potential near the threshold, suggesting the activation of inactivating currents (Hsu et al. 2018), though little is known about how these conductances affect synaptic summation *in vivo*. This raises questions about how these currents filter synaptic signals and determine synaptic summation modes.

Experimental results and computational modeling suggest that neurons process synaptic inputs via a two-stage integration process. In this process, synaptic inputs are integrated in individual dendritic branches, generating dendritic spikes that propagate to the axon, where global summation occurs (Katz, 2009; Polsky, 2004). Based on these findings, we developed a computational model of a single dendritic branch in which *in vivo*-like synaptic input activates voltage-dependent currents, and the corresponding synaptic potentials are integrated to generate dendritic spikes.

Understanding how ionic currents shape membrane temporal properties is essential for dendritic computation, influencing whether dendrites function as integrators or coincidence detectors. This paper explores the effects of A-type potassium, T-type calcium, and transient sodium currents on synaptic temporal summation *in vivo*-like conditions, focusing on their impact on voltage traces and summation properties (integrator vs. coincidence detector).

## Methods

Below, we introduce the single dendritic branch model, describe its embedded ionic currents, explain how the simulations were performed, and detail the mathematical analysis used to understand how these currents reduce/contract and amplify/prolong signals.

### Single dendritic branch model

We built a point-neuron model of a single dendritic branch with ionic currents, using the Hodgkin–Huxley formalism. The most general case of a Hodgkin-Huxley (HH) current involves two gating variables: *p* for activation and *q* for inactivation. This type of current is referred to as “inactivating” and acts as a break to close the channel (Hille, 2001). Each gating process has its own time constant, denoted as τ_act_ for activation and τ_inact_ for inactivation.

The current equation is

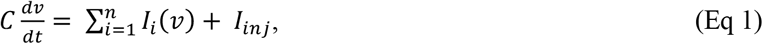

where *C* is the capacitance, *v* is the voltage, *I*_*i*_(*v*) is the voltage-dependent current, *n* is the number of ionic currents and *I*_*inj*_ is the injected current. In our paper, ionic currents will be represented using the HH formalism:

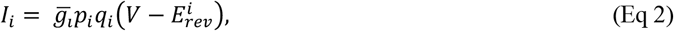

where 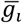 is the maximum conductance, 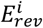 the reversal potential, and for (*x* = *p,q*) we have

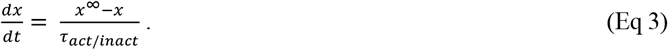

In Eq. 3, *x*^∞^ is the steady-state activation variable and it is represented by a sigmoid function following

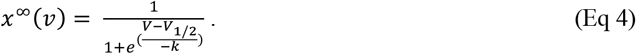

Where V_1/2_ and k are constants. Importantly, *p* = *p*(*v,t*) and *q* = *q*(*v,t*), i.e., activation and inactivation variables are dependent on voltage and time.

In this paper we only address inactivating currents activated by depolarization such as transient sodium, A-type potassium and T-type calcium. One important mathematical property of *p* and *q* in this situation that can be derived from the fact that they are activation and inactivation functions is that 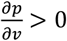 and 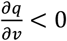, although p > 0 and q > 0. This behavior occurs for currents activated by depolarization, opposed to when the currents are activated by hyperpolarization 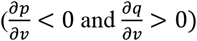.

Importantly, many inactivating potassium, sodium, and calcium currents follow Equation 2, while some variations of this equation exist where the activation or inactivation variables are raised to an exponent (such as 2, 3 or 4). Thus, the equation can be represented as 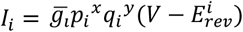. Although we will use this representation in our simulations, we will use the simple version as shown in Eq.2 for the analytical approximation.

We performed computer simulations using custom-made codes in Matlab. Three types of currents were investigated: A-type potassium, T-type calcium and transient sodium. Capacitance density was 1μF/cm^2^. Activation variables were defined by the following parameters: for potassium: *V*_1/2_ = -50 mV, *k* = 15, for calcium: *V*_1/2_ = -50 mV, *k* = 15, and for sodium: *V*_1/2_ = -30 mV, k = 5. Inactivation variables were defined by the following parameters: for potassium: *V*_1/2_ = -60, k = 15, for calcium: *V*_1/2_ = -60 mV, *k* = 15, and for sodium: *V*_1/2_ = -60 mV, k = 5. Activation kinetics was τ_act_ = 0.2, 0.5 and 0.05 ms for potassium, calcium and sodium respectively.

Inactivation kinetics was τ_inact_ = 5, 10 and 0.5 ms for potassium, calcium and sodium respectively. Conductance densities were expressed as mS/cm^2^ with values: *g*_K_ = 1, *g*_Ca_ = 0.1 and *g*_Na_ = 50. Leak conductance density was 0.3 mS/cm^2^. Reversal potential was *E*_rev_ = -80, 50 and 50 mV, for potassium, calcium and sodium respectively. Leak conductance reversal potentials was -80 mV.

The conductance values were chosen based on their ability to induce a sufficient depolarization or hyperpolarization of the resting membrane potential (approximately a 5 mV change) compared to the passive model (g_K_ = 10 mS/cm^2^, g_Ca_ = 0.1 mS/cm^2^). However, due to the explosive behavior of the sodium current—which can induce a rapid and uncontrollable depolarization beyond the firing threshold—we limited the sodium conductance to g_Na_ = 50 mS/cm^2^, resulting in a depolarization of approximately 1 mV (see Fig. 1). In some cases, we reduced the potassium conductance to 1 mS/cm^2^ to prevent complete silencing of spiking.

**Fig 1.**
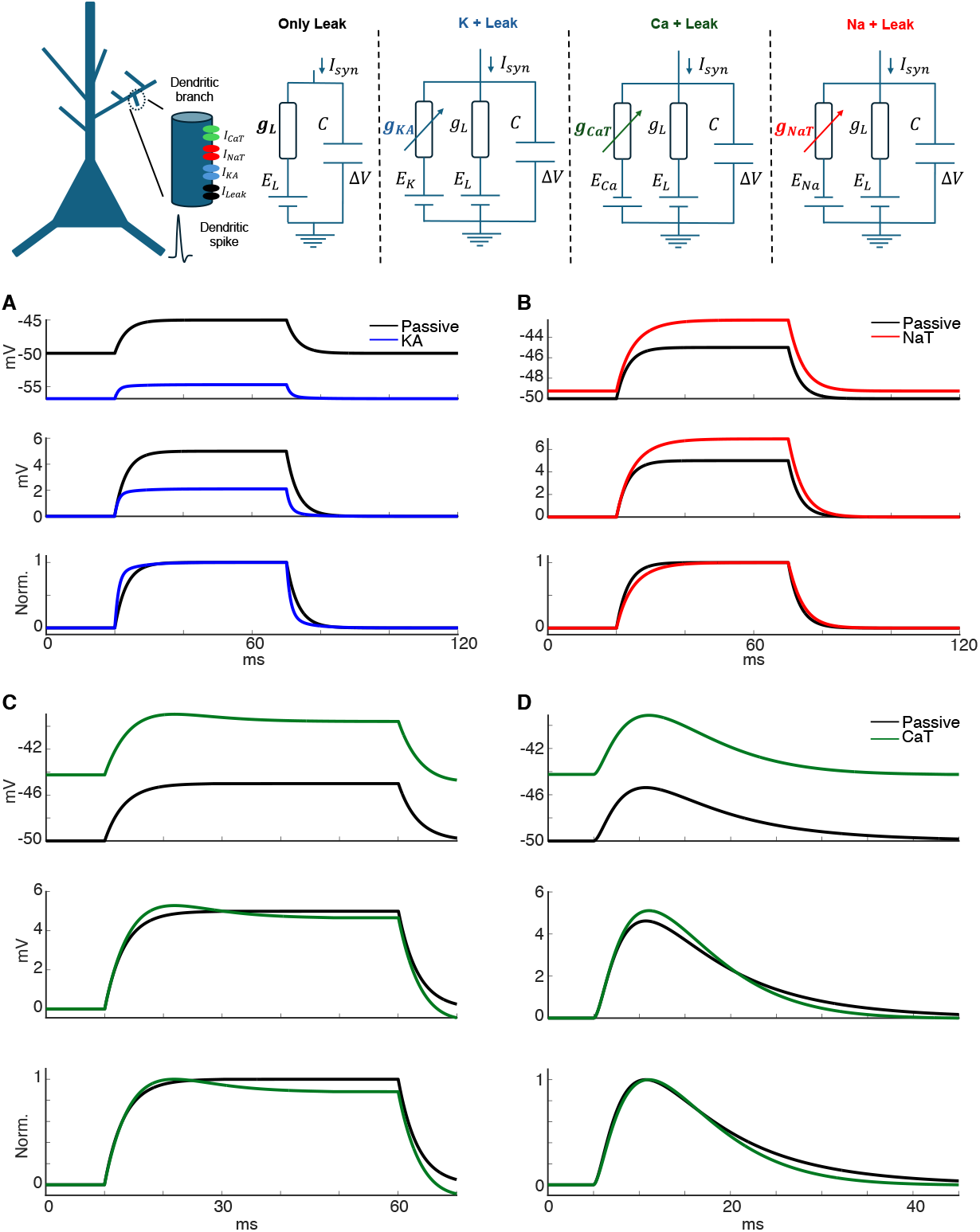
Membrane charging in the presence of the different ionic currents. Sketch at the top represents the four single dendritic branch models used in the simulations: Only leak (or passive, black), leak plus potassium current (blue), leak plus calcium current (green) and leak plus sodium current (red). In panels A, B, C and D, a current pulse of 50 ms was injected and the voltage change is displayed in the plots that are organized as: the original voltage trace of the relevant current plus leak model (coloured) and passive model (black) (upper), after baseline subtraction (middle), and after normalization (bottom). A. Voltage trace of the model with leak current plus *I*_KA_. B. Voltage trace of the model with leak current plus *I*_NaT_. C. Voltage trace of the model with leak current plus *I*_CaT_. D. An excitatory synaptic current was injected and the voltage changes are displayed in the plot that is organized as: the original voltage trace of the *I*_CaT_ plus leak current model (coloured) and passive model (black) (upper), after baseline subtraction (middle), and after normalization (bottom). In the simulations, the initial resting membrane potential is -50 mV. The conductance values were: g_KA_ = 10, g_CaT_ = 0.1 and g_NaT_ = 50, respectively. EPSCs were generated using biexponential events: *E*_rev_ = 0 mV, *g* = 0.04 nS. The time constants are τ_rise_ = 1 ms, τ_decay_ = 10 ms.

Models of A-type potassium, T-type calcium and transient sodium currents are a modified version from Engbers et al. 2012; Jaffe et al. 1994; Korngreen et al. 2000; Dougalis et al. 2017; and Hoffman et al. 1997. A-type potassium current equation is: 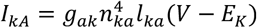. T-type calcium current equation is: 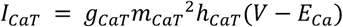. Transient sodium current equation is: 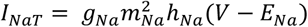. Leak current equation is: *I*_*L*_ = *g*_*L*_(*V* − *E*_*L*_).

In the next subsection, we describe how the simulations of these differential equations were performed.

### Simulations

Differential equations were solved using the Euler method and time step 0.05 ms. Synaptic background activity was generated of evenly distributed 200 excitatory postsynaptic currents (EPSCs) and 200 inhibitory postsynaptic currents (IPSCs) over time, to generate sparse or synchronous synaptic backgrounds as found *in vivo* (Waters et al., 2006, Harris and Thiele 2011). EPSCs and IPSCs were generated using biexponential events: *V*_Hold_ = -50 mV, *E*_AMPA_ = 0 mV, *E*_GABAA_ = -60 mV and *g* = 0.1 nS. The time constants are τ_rise/AMPA_ = 1 ms, τ_decay/AMPA_ = 10 ms, τ_rise/GABAA_ = 2 ms, τ_decay/ GABAA_ = 20 ms. For the synchronous case, we randomly removed 75% of the EPSCs and amplified the remaining ones by a factor of 4, with the goal of reproducing a highly synchronous set of inputs.

We simulated theta oscillations at *f* = 4 Hz and analyzed four cycles of voltage traces using the equation *I*_θ_ = *Asin*(2π*ft*), where *A* is the amplitude (5 μA/cm^2^) and *t* is time in ms. Thus, we injected in total the sum of the synaptic background activity with theta oscillations: *I*_*inj*_= *I*_*syn*_+ *I*_θ_, in other words, the *in vivo*-like state was generated as the combination of synaptic background activity with theta oscillations. We generated dendritic spikes using a leaky-integrate and fire model with resting potential -60 mV and spiking threshold -40 mV.

In the next subsection, we will describe our procedure to determine how reduction/contraction or amplification/prolongation occurs with the currents studied.

### Mathematical analysis of reducing/contracting and amplifying/prolonging currents

Here we used an analytical approach to determine the effect of *I*_KA_, *I*_CaT_, and *I*_NaT_ on the input resistance and membrane time constant. It is well known that τm is a function of *I*_i_:

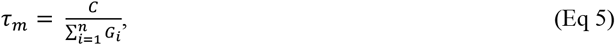

where the input conductance G_i_ can be obtained by differentiating Eq.2:

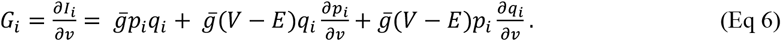

In this study all currents are inactivating. As shown in the results section (see Slow inactivation plays nearly no role on summation during *in vivo*-like states allowing for model reduction), all the inactivating currents produce similar voltage changes when it is assumed fast activation/inactivation kinetics. Under these circumstances, it is reasonable to assume τ_act_ = τ_inact_ → 0. Consequently, p and q become functions of voltage alone, denoted as p = p(V) and q = q(V). In this state, they lose their time dependence and are solely voltage dependent. Specifically, *p* = *p*^∞^(*v)* and *q* = *q*^∞^(*v)*. Therefore, when τ_act_ = τ_inact_ → 0, Eq 6 can be expressed as:

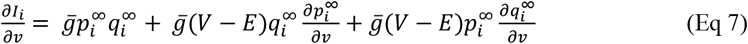

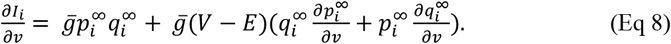

The term 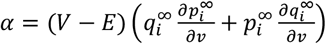 can assume positive or negative values. According to Eq. 5, when α > 0, τ_m_ decreases and voltage changes are reduced, whereas when α < 0, τ_m_ increases, and voltage changes are amplified. Currents with negative α are termed “amplifying/prolonging” as they lengthen τ_m_, while those with a positive α are termed “reducing/contracting”, as they shorten τ_m_ (Ceballos et al. 2017b).

The question arises: under what circumstances do the currents “amplify/prolong” or “reduce/contract”? To answer this, we need to investigate the conditions when α is positive or negative. We know that:

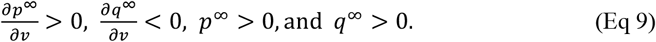

Therefore, 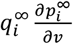 is positive and 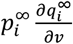 is negative. We also know that the membrane potential of dendrites typically ranges between -90 mV and +60 mV. The reversal potential for potassium (E_k_) is usually assumed to be around -90 mV, while the reversal potential for sodium and calcium (E_Na,Ca_) is assumed to be around +60 mV. Thus, for *I*_K_ ocurrs *V* − *E*_*5*_ > 0. Similarly, for *I*_CaT_ and *I*_NaT_, V-E_Na,Ca_ <0.

Substituting these into Eq. 7, we find that for I_K_ the first term is positive, the second is positive and the third is negative. Meanwhile, for *I*_CaT_ and *I*_NaT_, the second is negative and the third is positive. This demonstrates that activation and inactivation have opposite effects on τ_m_: one prolongs it while the other shortens it. This observation is consistent with the findings of Remme et al. (2011) and Richardson et al. (2003) that *I*_KA_ activation reduces, and inactivation amplifies the voltage scaling, and that *I*_KA_ activation shortens, and inactivation prolongs τ_m_. Meanwhile, the opposite effect occurs for *I*_CaT_ and *I*_NaT_.

A special case arises when inactivation is removed. This occurs for some voltage-dependent currents that exhibit either no inactivation or very slow inactivation (also known as nonactivating currents). This scenario can be simulated by setting q = 1. Therefore, we ask: when can 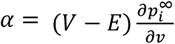 be positive or negative, i.e., when does it contract or prolong τ_m_, after inactivation is removed? We can easily deduce that, 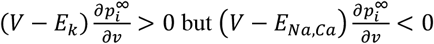. This suggests that when removing the inactivation from *I*_KA_, α is positive i.e., it is always a “contracting” current, meanwhile for *I*_CaT_ and *I*_NaT_, α is negative i.e., it is always a “prolonging” current.

## Results

### Consistent membrane charging effects in the presence of potassium and sodium currents, but mixed response effects in the presence of calcium current

We begin our paper testing the effect of inactivating currents on membrane time constant (τ_m)_. For that, we built a single compartment dendritic model. We focused on the most relevant inactivating currents expressed in pyramidal neurons dendrites and activated near the spiking threshold: A-type potassium (*I*_KA_), T-type calcium (*I*_CaT_), and transient sodium (*I*_NaT_) (see Methods). A depolarizing pulse of current was applied, and both exponential rise and decay followed by this pulse were analyzed. Fig. 1A shows that *I*_KA_ hyperpolarizes the resting membrane potential, reduces the amplitude of the response, and accelerates the capacitor’s charge. In contrast, *I*_NaT_ amplifies the response amplitude and slows down the capacitor’s charge but has no effect on the resting membrane potential due to its activation at higher membrane potentials (Fig 1B).

Opposed to this pattern, the effects of *I*_CaT_ were mixed. *I*_CaT_ depolarizes the resting membrane potential. However, there is only amplification of the voltage change at the onset of the current pulse, while there is a reduction of the voltage change at the end of the pulse due to slow inactivation kinetics (Fig 1C). We also observed a clear phase delay during the rising phase of the EPSP (excitatory postsynaptic potential), but the decay phase showed mixed behavior, starting with a phase delay in one direction and eventually reversing its direction (Fig 1D). We believe this mixed behavior in amplification/reduction of the voltage change and lengthening/shortening of the voltage provides insight into the inconsistent reports found in the literature about *I*_CaT_ (Stuart et al. 1995; González-Burgos et al. 2001; Andreasen et al. 1999; Hsu et al. 2018; Seong et al. 2014, Fukaya et al. 2018; Gillessen et al. 1997, Prescott et al. 2005, Liu et al. 2008; Connelly et al. 2016; Urban et al. 1998; Evans et al. 2017; Deleuze et al. 2012).

Moreover, we further investigated the combined effects of multiple ionic currents (*I*_KA_, *I*_CaT_ and *I*_NaT_) by measuring the difference in resting membrane potentials (ΔRMP), scaling, and the membrane time constant (τ_m_). In the heatmaps presented in Fig. S1, two conductance values were varied while one was fixed, allowing us to assess how the combination of these currents influences these measures. ΔRMP was calculated as the difference between the RMPs of the two traces, while scaling was determined as the ratio of the voltages (V_2_/V_1_). The membrane time constant was obtained by fitting the rising phase with a single exponential function. The results from this supplementary figure demonstrate a strong correlation between scaling and τ_m_, which is expected based on the mathematical relationship *τ*_*A*_ = *R*_*input*_ × *C*_*A*_, where scaling is analogous to the traditional input resistance. Furthermore, in the presence of the other currents, *I*_KA_ decreases τ_m_, whereas *I*_CaT_ and *I*_NaT_ increase τ_m_. Notably, *I*_CaT_ has a mild effect on τ_m_ but a strong effect on the resting membrane potential.

### Voltage changes scaling monotonically during theta oscillations

The previous section studied the effect of the inactivating currents on the temporal properties of a dendritic model. The model was stimulated using simple current pulses and simulated EPSPs. Here, we wondered whether the previous results also apply under *in vivo-*like conditions. Here, we investigated the effects of the inactivating currents on scaling voltage traces from *in vivo*-like conditions. We injected synaptic currents, consisting of evenly distributed EPSCs and IPSCs over time, to generate sparse synaptic backgrounds as found *in vivo* (Waters et al. 2006). To add realistic background to the synaptic currents, we simulated theta oscillations at 4 Hz and analyzed four cycles of voltage traces (Fig. 2; cycles identified by different colors). Our results show that *I*_KA_ causes a negative shift of the voltage changes (Fig 2A), whereas *I*_CaT_ and *I*_NaT_ cause a positive shift (Fig 2C, 2E). The scaling from above -60 mV is linearly correlated with the membrane voltage for *I*_KA_ and *I*_NaT_ (Fig 2B, 2F) but no clear tendency is found for *I*_CaT_, only high variability (Fig 2D). The data presented in Fig. 2 reveal that *I*_KA_ induces a negative shift with scaling at approximately 90% at the peak of a cycle, whereas *I*_NaT_ causes a positive shift with scaling at around 115%. When translated to voltage, this corresponds to a shift of approximately 6 mV. Given that we are operating in a regime near the action potential threshold and considering the context of an *in vivo*-driven neuron, this shift can be decisive in determining whether a spike occurs. These findings indicate that *I*_KA_ and *I*_NaT_ can modulate spiking activity in an oscillatory state, highlighting the significance of these currents in shaping neuronal excitability.

**Fig 2.**
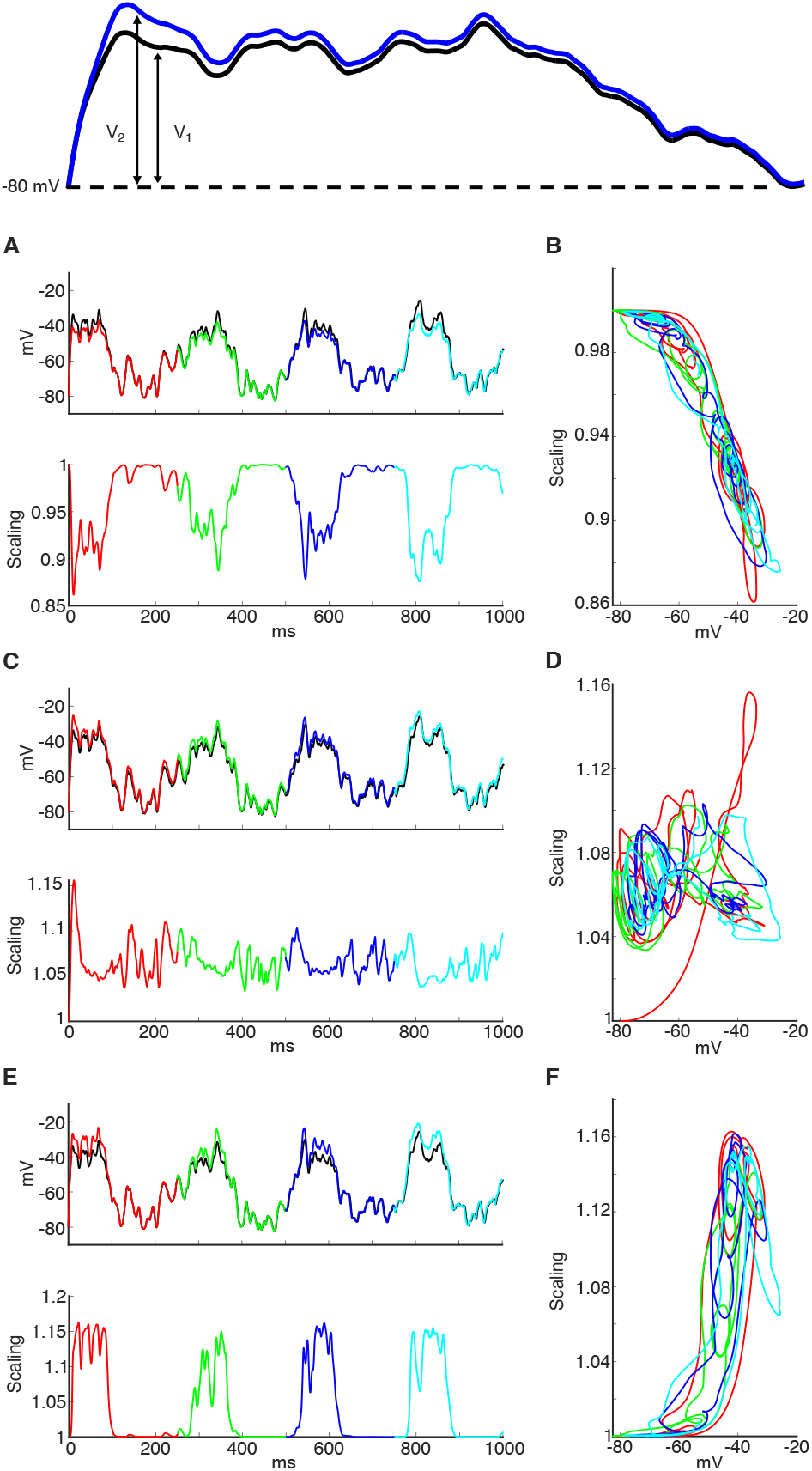
Sketch of voltage traces and the measurement of the relative voltage used to calculate the scaling. Voltage changes for dendritic ionic currents under *in vivo*-like conditions. Dendritic models were stimulated by injection of a 4 Hz theta oscillations plus synaptic background made of randomly distributed excitatory and inhibitory synaptic currents. The plot displays the voltage traces of the passive dendrite (black solid line). Red/Green/Blue/Cyan represent first to fourth cycle of the theta sequence in the dendrite models with leak current plus the relevant ionic current. A. Voltage traces and scaling of dendrite expressing only leak current plus *I*_KA_. B. phase plot of nonlinear scaling vs membrane potential. C. Voltage traces and scaling of dendrite expressing only leak current plus *I*_CaT_. D. phase plot of nonlinear scaling vs membrane potential. E. Voltage traces and scaling of dendrite expressing only leak current plus *I*_NaT_. F. phase plot of nonlinear scaling vs membrane potential. Scaling was defined as (V_2_ + 90) / (V_1_ + 90), where V_2_ is the voltage trace of the neuron model with both leak and relevant currents, and V_1_ is the voltage trace with only the leak current. Since the lowest simulated voltage is -80 mV, we added 90 mV to ensure all values remain positive. Phase plots are showing the voltage-dependence of the scaling. This is an analogous measurement of the conventional way to investigate the amplification/shrinking of EPSPs by voltage-dependent currents found in in vitro studies. The conductance values were: g_KA_ = 1, g_CaT_ = 0.1 and g_NaT_ = 50, respectively.

Fig 3 shows the nonlinear scaling for input frequency oscillations from 1 to 30 Hz. The set of Figs 3A-C demonstrate a clear difference in the voltage scaling when comparing the ionic currents. More specifically, we observed a consistent decrease for *I*_KA_ across depolarizing voltages and a consistent increase for *I*_NaT_ across depolarizing voltages (see heatmaps in Fig 3D and Fig 3F). In contrast, *I*_CaT_ showed nonmonotonic behavior of the scaling for low frequencies, but monotonic for high frequencies (Fig 3E). Then we evaluated the behavior of the scaling for different conductance values at 4 Hz. We observed a consistent decrease for I_KA_ across depolarizing voltages and a consistent increase for *I*_NaT_ across depolarizing voltages (Fig 3G and Fig 3I). In contrast, *I*_CaT_ showed similar values across voltage (horizontal lines, Fig 3H).

**Fig 3.**
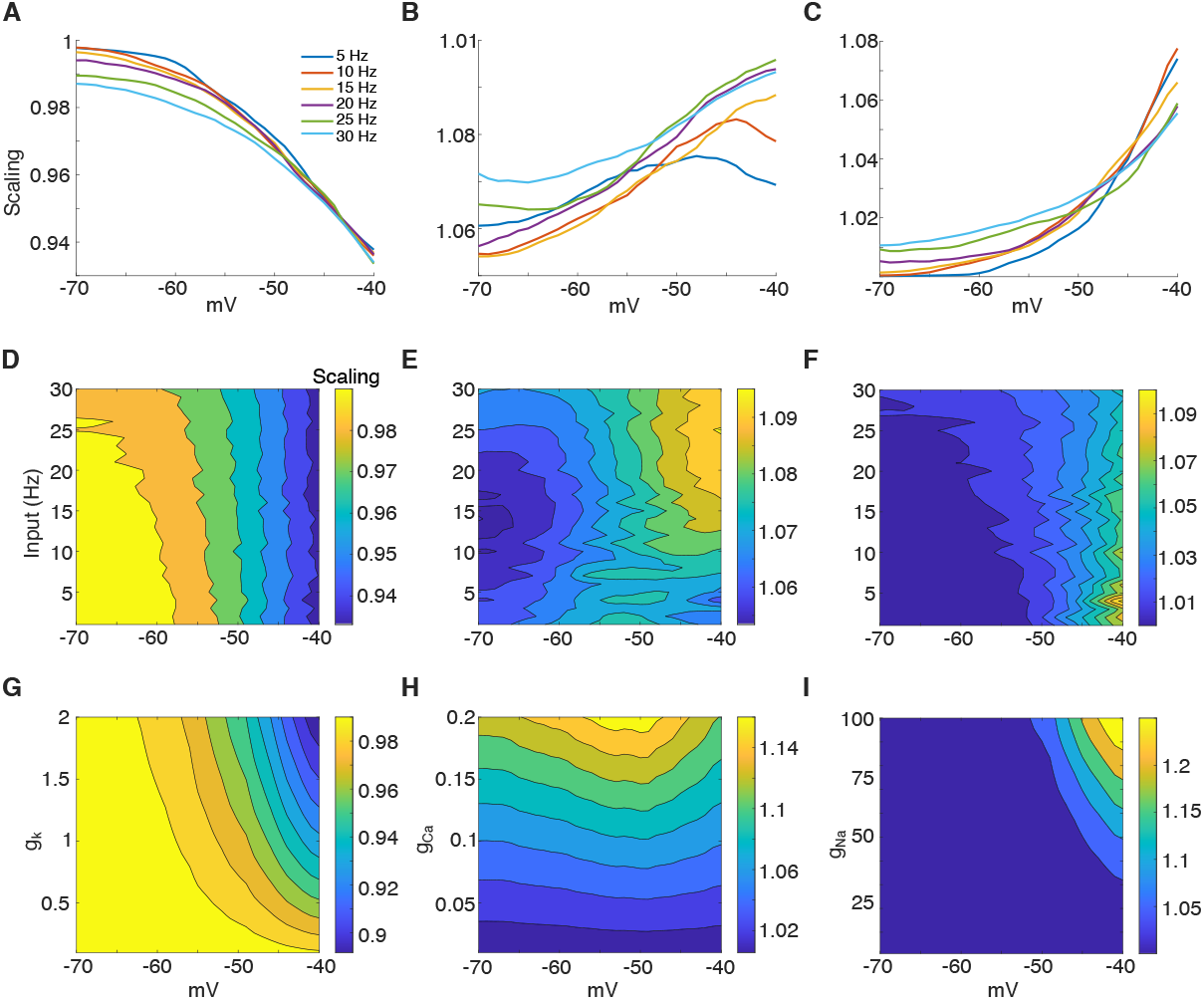
Frequency and ionic effect on voltage scaling. The nonlinear scaling is computed when input frequency oscillations change range from 1 to 30 Hz. A. Curves for added *I*_KA_. B. Curves for added *I*_CaT_. C. Curves for added *I*_NaT_. D. Heatmap of the nonlinear scaling for voltages from -70 to -40 mV vs. input frequency oscillations change from 1 to 30 Hz for *I*_KA_. E. Same as in D but for *I*_CaT_. F. Same as in D but for *I*_NaT_. G. Heatmap of nonlinear scaling for voltages from -70 to -40 mV vs. *I*_KA_ conductance. g_CaT_ = 0.1 and g_NaT_ = 50. H. Same as in G but for *I*_CaT_ conductance. g_KA_ = 1 and g_NaT_ = 50. I. Same as in G but for *I*_NaT_ conductance. g_KA_ = 1 and g_CaT_ = 0.1. Frequency was fixed at 4 Hz for G, H and I plots. Scaling was defined as (V_2_ + 90) / (V_1_ + 90), where V_2_ is the voltage trace of the neuron model with both leak and relevant currents, and V_1_ is the voltage trace with only the leak current. Since the lowest simulated voltage is -80 mV, we added 90 mV to ensure all values remain positive. The conductance values were: g_KA_ = 1, g_CaT_ = 0.1 and g_NaT_ = 50, respectively.

These results suggest that most of the knowledge from Fig 1 can be transferred to the *in vivo*-like states except for *I*_CaT_, where its ionic currents scale the voltage differently in a frequency-dependent manner.

### Changes in activation are linearly correlated with changes in inactivation for T-type calcium and A-type potassium currents

We previously observed the effect of the inactivating currents on the scaling of the voltage. *I*_KA_ and *I*_NaT_ showed linear correlation with voltage, but no consistent tendency was observed for *I*_CaT_. In this section, to investigate the biophysical mechanisms underlying these discrepancies we present phase plots of activation versus inactivation variables for theta oscillations (Fig. 4). This analysis demonstrated a clear negative correlation, where increases in activation correspond with decreases in inactivation (Fig 4G, 4H, 4J and 4K). Interestingly, the window current properties of *I*_NaT_ are markedly different than *I*_KA_ and *I*_CaT_ and produced a highly nonlinear correlation between activation and inactivation variables, where large values of one variable correlate with small values of the other (Fig 4I). We verified that this relationship is well-fitted by a power law function (Fig 4L).

**Fig 4.**
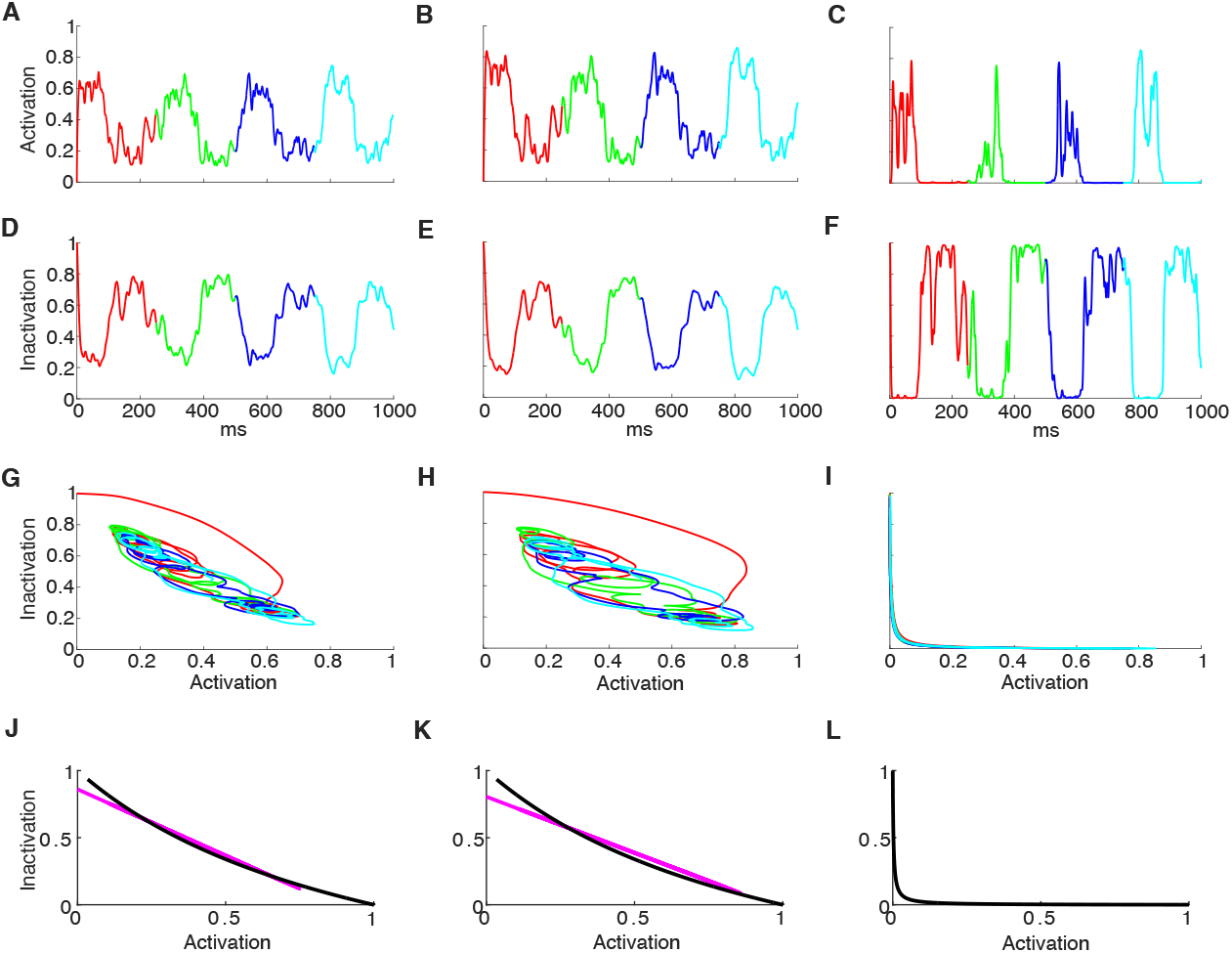
Activation vs. inactivation functional relationships. A-F. The curves represent activation and inactivation variables under 4 Hz theta oscillations. Red/Green/Blue/Cyan represent first to fourth cycle of the theta sequence with the added ionic current: activation for A. *I*_KA_. B. *I*_CaT_. C. *I*_NaT_; inactivation for D. *I*_KA_. E. *I*_CaT_. F. *I*_NaT_. G-I: Phase plots of activation vs inactivation for G. *I*_KA_. H. *I*_CaT_. I. *I*_NaT_. Notice the filtering of the inactivation variable due to the slow inactivation kinetics of *I*_CaT_ and *I*_KA_. J-L: Linear fitting of activation vs. inactivation phase plots for J. *I*_KA_. K. *I*_CaT_ (magenta) and power-law fitting for L. *I*_NaT_. Trajectories when τ_act_ = τ_inact_ = 0 (black). The conductance values were: g_KA_ = 1, g_CaT_ = 0.1 and g_NaT_ = 50, respectively.

Analysis of the activation vs. inactivation phase plots showed that fitting trajectories of *I*_KA_ and *I*_CaT_ by a linear regression returns a curve very similar to the trajectories when τ_act_ = τ_inact_ = 0 (Fig 4J and 4K). This allows us to approximate the functional relationship between activation or inactivation. Therefore, we use the linear regression to substitute the inactivation variable by a linear function dependent on the activation variable: q_Inact_ = - p_Act_ + 0.85 for *I*_KA_ and, q_Inact_ = - 0.84p_Act_ + 0.8 for *I*_CaT_. In contrast, the trajectories for *I*_NaT_ are highly nonlinear and, as explained above, were fitted using a power law function with 4/3 exponent 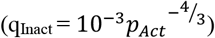 (Fig 4I and 4L). This functional behavior is the result of the way activation and inactivation curves interact within the window current.

Our results showed similar activation/inactivation linear trajectories for *I*_KA_ and *I*_CaT_, although these two currents differ in their inactivation kinetics (e.g. τ_inact_ = 5 ms for *I*_KA_ but 10 ms for *I*_CaT_). This suggests that under *in vivo*-like conditions, voltage changes are robust to inactivation kinetics.

### Slow inactivation plays nearly no role on summation during *in vivo*-like states

Based on the results above, we interrogate whether *in vivo*-like states require kinetics. Therefore, we investigated whether we could get similar voltage traces when using the actual kinetics vs. when using fast kinetics, i.e. τ_act_ = τ_inact_ = 0.1 ms. Surprisingly, for all simulated currents we obtained traces that were similar with average differences less than 0.6 mV for voltages below spiking threshold (V = -40 mV; Fig 5D). These results suggest that the system of equations can be reduced when assuming τ_act_ = τ_inact_ = 0, hinting that the dynamics provided by Eq 3 is no longer necessary for *in vivo*-like conditions. Suprathreshold spiking activity also displayed similar results (Fig S2).

**Fig 5.**
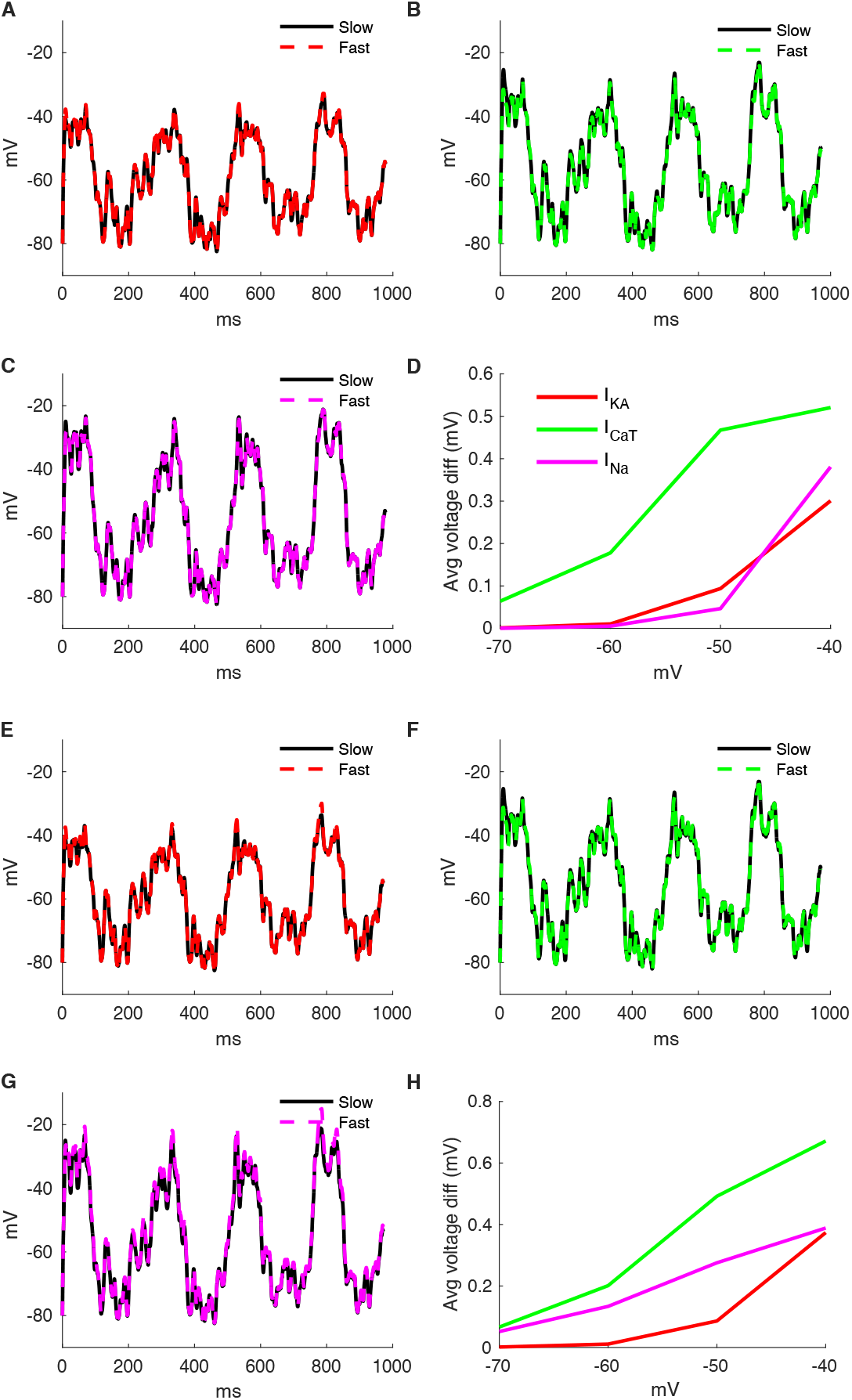
Comparison of traces using actual kinetics vs. fast kinetics (i.e. τ_act_ = τ_inact_ = 0.1 ms). A. *I*_KA_. B. *I*_CaT_. C. *I*_NaT_. D. Average voltage difference between the two tested conditions (*I*_KA_ in red, *I*_CaT_ in green, *I*_NaT_ in magenta). Comparison of traces using actual activation/inactivation variables and kinetics vs. fast kinetics (i.e. τ_act_ = τ_inact_ = 0 ms) and after linearizing inactivation variables using activation variable for *I*_KA_ and *I*_CaT_ and power law approximation for *I*_NaT_. E. *I*_KA_. F. *I*_CaT_. G. *I*_NaT_. H. Average voltage difference between the two tested conditions (*I*_KA_ in red, *I*_CaT_ in green, *I*_NaT_ in magenta). The conductance values were: g_KA_ = 1, g_CaT_ = 0.1 and g_NaT_ = 50, respectively.

Further, and following the relationships from the previous section, we can also reduce the inactivation variable using the linear relation obtained from the linear regression above for *I*_KA_ and *I*_CaT_ and power law approximation for *I*_NaT_ inactivation. As a result of all the reductions, we can express the behavior of each current as dependent on only a single activation variable. Therefore, we use the linear regression to substitute the inactivation variable by a linear function dependent on the activation variable for *I*_KA_ and *I*_CaT_ and power law approximation for *I*_NaT_. We also consider τ_act_ = τ_inact_ = 0. Fig 5H demonstrates that this approximation can obtain similar voltage traces with average values below 0.7 mV for all currents.

The results from this section allowed us to reduce the dendrite system of equations to a single variable that resolves the activation variable only. Our approximation is akin to the Morris-Lecar reduction (Izhikevich 2007). However, in our case, the assumption of instantaneous kinetics allows us to avoid coupling differential equation.

In the next set of experiments, we wondered about the dependency of our results with the input frequency. Fig 6 shows the voltage difference between the model with slow kinetics vs. instantaneous for input frequency oscillations from 1 to 30 Hz. We observed a consistent increase for all currents across depolarizing voltages, with values below 0.7 mV for *I*_KA_, 1.2 mV for *I*_CaT_ and 0.35 mV for *I*_NaT_ (Fig 6D, 6E and 6F). Then we evaluated the voltage difference between the model with slow kinetics vs. instantaneous kinetics for different conductance values with input frequency fixed at 4 Hz. We observed a consistent increase for all currents across depolarizing voltages, with lowest values at conductance close to g_KA_ = 1, g_CaT_ = 0.1 and g_NaT_ = 50 (Fig 6G, 6H and 6I).

**Fig 6.**
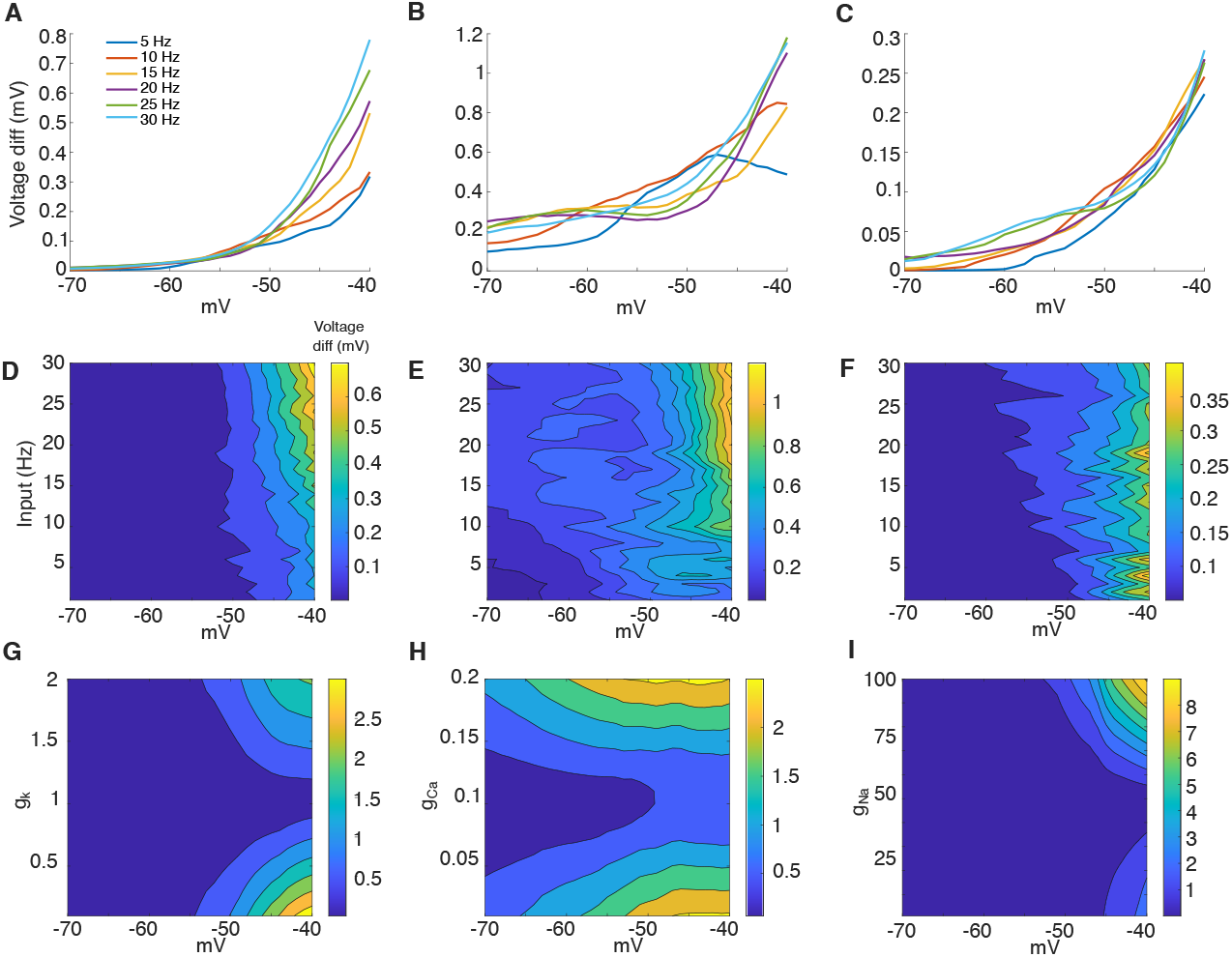
Voltage difference between the model with slow kinetics vs. instantaneous, when input frequency oscillations change from 1 to 30 Hz. A. *I*_KA_. B. *I*_CaT_. C. *I*_NaT_. D. Heatmap of voltage difference between the model with slow kinetics vs instantaneous for voltages from -70 to -40 mV vs. input frequency oscillations change from 1 to 30 Hz for *I*_KA_. E. Same as in D but for *I*_CaT_. F. Same as in D but for *I*_NaT_. G. Heatmap of the voltage difference between the model with slow kinetics vs instantaneous for voltages from -70 to -40 mV vs. I_KA_ conductance. g_CaT_ = 0.1 and g_NaT_ = 50. H. Same as in D but for *I*_CaT_ conductance. g_KA_ = 1 and g_NaT_ = 50. I. Same as in D but for *I*_NaT_ conductance. g_KA_ = 1 and g_CaT_ = 0.1. The conductance values were: g_KA_ = 1, g_CaT_ = 0.1 and g_NaT_ = 50, respectively.

These results suggest that the voltage differences between the model with slow kinetics and fast kinetics is independent of the frequency input, but dependent on the voltage. As to the latter, as the depolarization became stronger, the voltage difference identified grew greater, showcasing that the kinetics becomes more prominent as the voltage gets closer to the threshold of the neuron.

### Inactivation opposes activation effects on voltage scaling and membrane time constant

We know from above that activation and inactivation have negative correlation, but it is still unknown how this correlation determines the voltage scaling and temporal properties of the voltage traces in the presence of *I*_KA_, *I*_CaT_ and *I*_NaT_. On the other hand, it is well known that scaling is inversely related to the input conductance and that the temporal properties are related to the membrane time constant. We first used an analytical approach to determine the opposed effect of the inactivation on voltage scaling and temporal properties compared to activation. When comparing the input conductance for inactivating current (as shown in Eq 8) vs non-inactivating current (as shown in Eq 10), we know from Eq 9 that 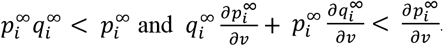. Thus, input conductance of inactivating currents is always smaller compared to the non-inactivating case, suggesting that the effect of inactivation is always to oppose to the activation.

To test the opposite effects of activation and inactivation, we investigated the temporal summation of a train of 3 EPSPs separated by 10 ms. We tested the effect of the inactivating current and their respective non-inactivating version (Fig 7). These conditions are represented by the following equations: current with inactivation 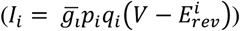, and current without inactivation or non-inactivating 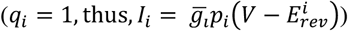. The plots show that noninactivating currents enhance the effect compared to their inactivating counterpart. These results suggest that inactivation counteracts the effect of the activation, reducing its effect.

**Fig 7.**
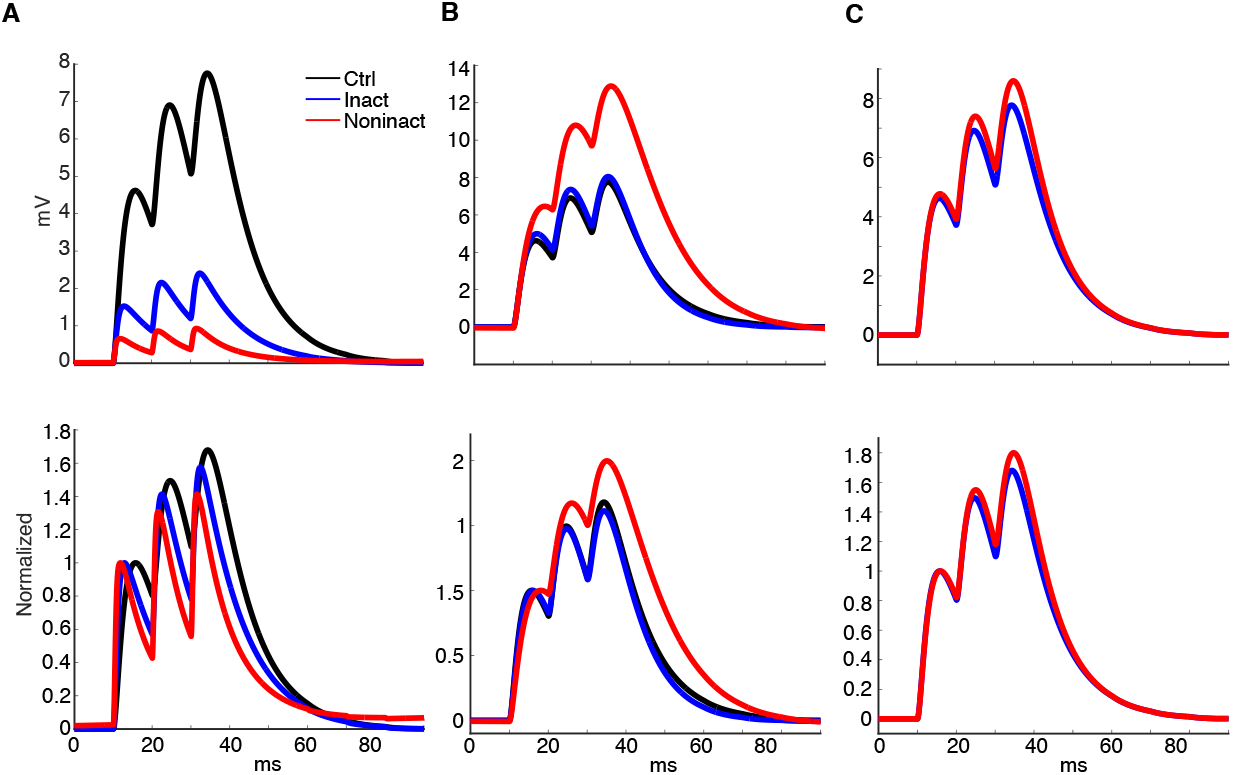
Inactivation and activation effects on summation. The voltage traces show summation for a train of 3 excitatory synaptic potentials at 100 Hz. A. Model with *I*_KA_. B. Model with *I*_CaT_. C. Model with *I*_NaT_. Initial resting membrane potential is -50 mV. Top: after baseline subtraction, bottom: after normalization. Three conditions were analyzed: only leak current (black), leak + inactivating current (blue), leak + noninactivating current (red). These conditions are represented by the following equations: current with inactivation 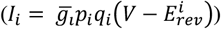, and current without inactivation or non-inactivating 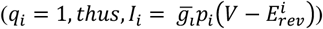. EPSCs were generated using biexponential events: *E*_rev_ = 0 mV, *g* = 0.04 nS. The time constants are τ_rise_ = 1 ms, τ_decay_ = 10 ms. The conductance values were: g_KA_ = 10, g_CaT_ = 0.1 and g_NaT_ = 0.5, respectively.

While in this section we showed that inactivation opposes activation effects on voltage scaling and membrane time constant with analytical derivations, together with previous sections we can conclude that *I*_KA_ activation shortens τ_m_ whereas inactivation prolongs it (Fig 7A). Meanwhile, an opposite effect occurs for *I*_CaT_ and *I*_NaT_ (Fig 7B and 7C). Together, these observations have profound implications for dendritic computation, in particular to their temporal summation properties. In the next section, we will show how our results are useful to study synchrony- vs. rate-code.

### A-type potassium current enhances synchrony-code but T-type calcium and transient sodium currents enhance rate-code

Temporal summation of synaptic potentials can be used to identify integrators or coincidence detector neurons. Integrators, characterized by a long τ_m_, allow summation of synaptic potentials even when they occur far apart in time. In contrast, coincidence detectors, with a short τ_m_, integrate synaptic potentials only when they occur closely together in time. Previous studies have shown the association between high expression of potassium currents and coincidence detection mode, and high expression of calcium/sodium currents and integration mode (Mathews et al. 2010; Branco et al. 2016). We tested this hypothesis using an analytical approach to demonstrate that for instantaneous inactivating currents, potassium currents such as *I*_KA_ always decrease the input resistance and shorten the membrane time constant, meanwhile calcium/sodium currents such as *I*_CaT_, and *I*_NaT_ always increase the input resistance and prolong the membrane time constant (see Methods, section Mathematical analysis of reducing/contracting and amplifying/prolonging currents).

Moreover, coincidence detection and integration on the level of dendritic compartments can be seem translated to synchrony-code and rate-code on the level of network computations, with coincidence detectors being more likely to fire with synchronized inputs and poorly integrate sparse inputs. In contrast, integrators can summate sparse inputs to generate dendritic spikes but respond less to coincident inputs compared to coincidence detectors. Thus, a natural question we investigate in this section is how the modulation of the *I*_KA_, *I*_CaT_ and *I*_NaT_ affect the synchrony vs. rate-code.

Accordingly, we randomly generated 100 sparse and 100 synchronous input currents and used them to generate 100 voltage traces also stimulated by theta input (4 Hz). The sparse and synchronous input were developed to create distinction between these codes (synchrony- vs. rate-code). Both sparse and synchronous inputs had the same amount of 200 EPSCs and 200 IPSCs randomly distributed in the course of a simulation. For the synchronous case, we removed randomly 75% of the EPSCs and the remaining were amplified by a factor of 4 with the rationale of reproducing a highly synchronous set of inputs.

While a consistent decrease in firing occurs in the presence of the *I*_KA_ (compare Fig 8A to B), an increase in firing occurs for *I*_CaT_ and *I*_NaT_ (compare Fig 8A to C and D). This aligns with the shrinkage and amplification of voltage changes seen above. When all three currents were present, firing rates remained unchanged (not shown), indicating that the effects of *I*_KA_ were counteracted by *I*_CaT_ and *I*_NaT_ as expected. The figure demonstrates the ability of ionic currents to transform a neuron into a coincident detector that is more likely to process synchronized inputs and less likely to integrate sparse inputs. This is the case for *I*_KA_, as it more effectively processes synchronous inputs (see histograms in Fig. 8F). The other currents do not exhibit this ability, processing synchronous and sparse inputs equally.

**Fig 8.**
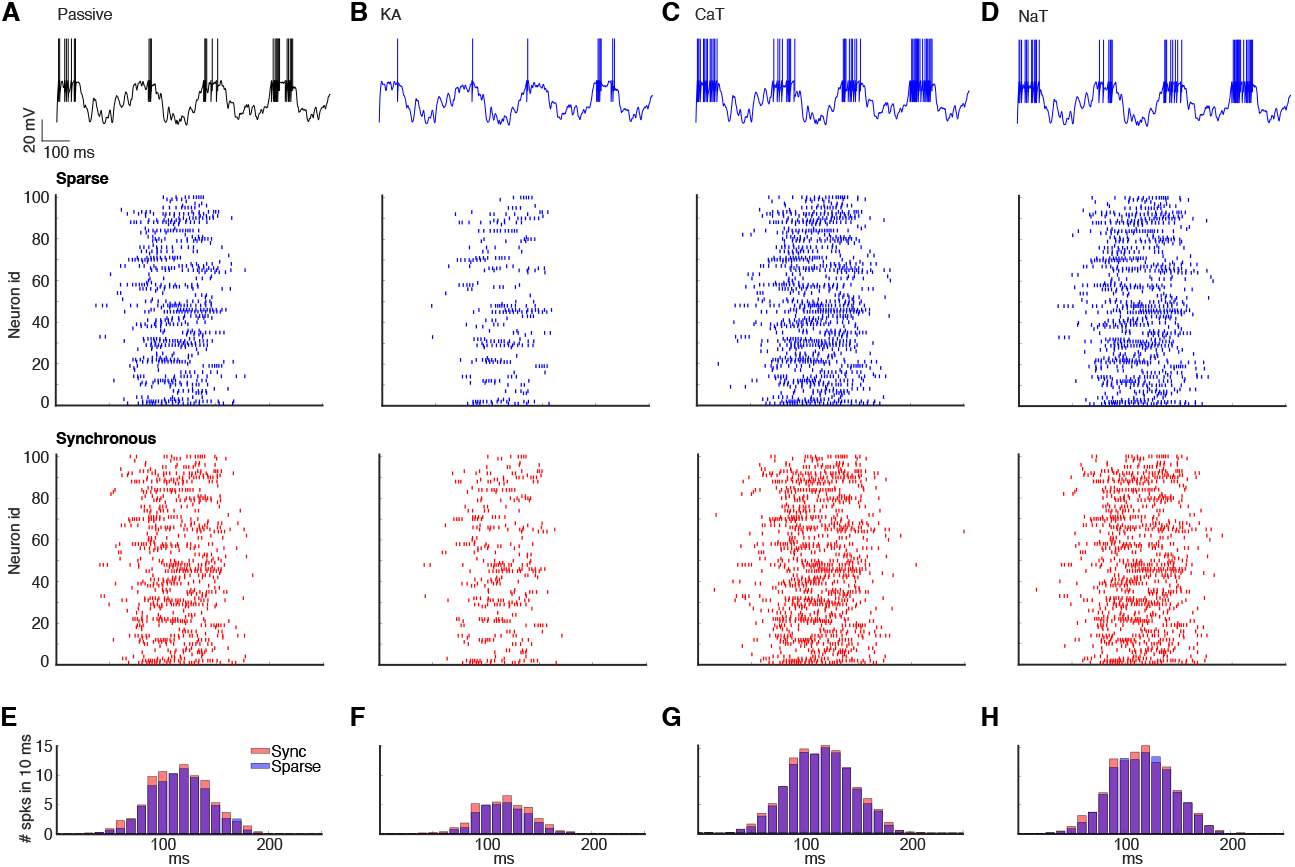
Synchrony- vs. rate-code and ionic currents dependency. Every column in the figure contains an exemplary voltage trace followed by raster plots of the sparse and synchronous inputs, as well as a post-stimulus time histogram (PSTH). Firing pattern under 4 Hz theta oscillations in a dendrite expressing. A. only leak current. B. dendrite expressing only leak current plus *I*_KA_. C. neuron expressing only leak current plus *I*_CaT_. D. dendrite expressing only leak current plus *I*_NaT_. (E, F, G, H). The figure demonstrates the ability of ionic currents to transform a neuron into coincident detector which is more likely to process synchronized inputs and less integrate sparse inputs, which is the case of *I*_KA_. The conductance values were: g_KA_ = 1, g_CaT_ = 0.1 and g_NaT_ = 50, respectively.

To further quantify our results in Fig. 8, we require a metric that can determine the preferred synaptic summation mode (coincidence vs. integration). Thus, we calculated the relative change in spiking frequency of a dendrite when presented with sparse versus synchronized synaptic inputs 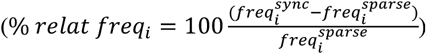, where 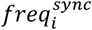 is the spiking frequency when stimulated with synchronized input and *freq*^*sparse*^ is the spiking frequency when stimulated with sparse input. *i* is the corresponding relevant current (*I*_KA_, *I*_CaT_ or *I*_NaT_). We expect that coincidence detectors will show an increase in relative spiking frequency, while integrators will show a decrease. We tested this new metric in a single passive compartment dendrite model by changing the conductance value or capacitance density (Fig S3). The heatmap shows the percentage of relative frequency change when dendrites were stimulated using sparse vs synchrony input. We used this index (% relative frequency) as a measure of the strength of the preferred neural code mode employed by the dendrite. For instance, % relative frequency of 100% suggests the dendrite acts as a perfect coincident detector, meanwhile 0% suggests the dendrite acts as a perfect integrator. According to the values obtained from the index, we found that the dendrite would act closer to a perfect integrator when g_Leak_ was low or when capacitance density was high, both conditions consistent with long τ_m_.

As shown in Fig 8, synchronous input increases firing rate. The relative increase in frequency is highest at the onset of the rise and decay of the theta cycle and lowest at the peak (Fig 8E, 8F, 8G and 8H). The exception is the potassium current where no large relative increase was observed at the decay (Fig 8F). We also calculated the relative difference of frequency between sparse vs synchronous stimulation (Fig 8E, 8F, 8G and 8H). We found a significant increase in the presence of *I*_KA_ relative to the passive dendrite (17.44 ± 1.38 % for passive, 38.86 ± 4.65 % for *I*_KA_, p <0.001). We also found a significant decrease in the presence of *I*_CaT_ and *I*_NaT_ relative to the passive dendrite (9.77 ± 0.81 % for I_CaT_, 10.53 ± 0.8 % for *I*_NaT_, p <0.001). Also, *I*_KA_ was significantly higher than *I*_CaT_ and *I*_NaT_ (p<0.001). Finally, *I*_CaT_ and *I*_NaT_ were not significantly different (p = 0.5). These results suggest that *I*_KA_ enhances synchrony-code, while *I*_CaT_ and *I*_NaT_ enhance rate-code.

We then expand our analysis to investigate the tuning range of specific neural code modes and how they depend on the conductance values of *I*_KA_, *I*_CaT_ and *I*_NaT_. We use the same dendritic model used above, but now we added all 3 currents to the model, while changing their conductance values selectively. According to the values obtained from the index, we found that the dendrite would act closer to a perfect integrator when g_KA_ was low (Fig 9A) or when g_CaT_/g_NaT_ were high (Fig 9H, I). Under these conditions, the variability of the other conductances did not affect much the index value (the heatmaps are blueish). These results suggest *I*_KA_ is responsible for changing the dendrite to a more coincidence detection mode meanwhile *I*_CaT_ and *I*_NaT_ take the dendrite closer to an integration mode. This can also be observed for high values of g_KA_ (see Fig 9G) or when g_CaT_/g_NaT_ is low (Fig 9B, C). Under these conditions the variability of the other conductances play a significant role that affects the index value (the heatmaps change from dark blue to light yellow).

**Fig 9.**
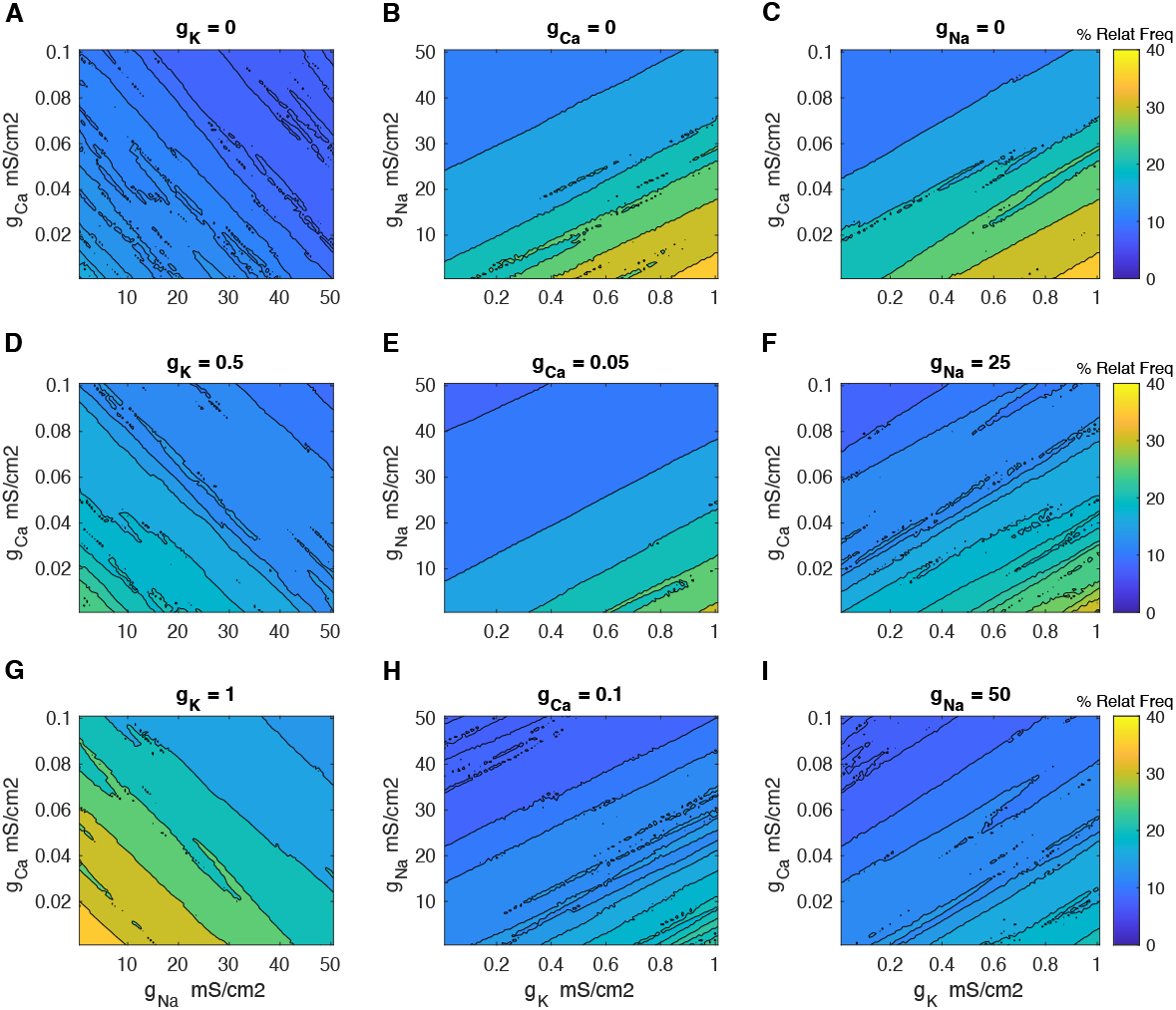
Heatmaps of % relative frequency (% relat freq) computed as difference between the sparse versus synchronized synaptic inputs spanned over different conductance values (see axis). Firing pattern under 4 Hz theta oscillations stimulation. A. *g*_NaT_ vs *g*_CaT_ when g_K_ = 0. B. *g*_KA_ vs *g*_NaT_ when g_Ca_ = 0. C. *g*_KA_ vs *g*_CaT_ when g_K_ = 0. D. *g*_NaT_ vs *g*_CaT_ when g_K_ = 0.5. E. *g*_KA_ vs *g*_NaT_ when g_Ca_ = 0.05. F. *g*_KA_ vs *g*_CaT_ when g_Na_ = 25. G. *g*_NaT_ vs *g*_CaT_ when g_K_ = 1. H. *g*_KA_ vs *g*_NaT_ when g_Ca_ = 0.1. I. *g*_KA_ vs *g*_CaT_ when g_Na_ = 50. Notice that the dendrite would act closer to a perfect integrator when g_KA_ was low or when g_CaT_/g_NaT_ were high, suggesting that *I*_KA_ is responsible for changing the dendrite to a more coincidence detection mode meanwhile *I*_CaT_ and *I*_NaT_ take the dendrite closer to an integration mode. This can also be observed for high values of g_KA_ or when g_CaT_/g_NaT_ is low. The % relative frequency is an indicator of whether the dendrite acts as an integrator of coincidence detector (100% for perfect coincident detector and 0% for perfect integrator). 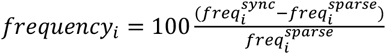, where 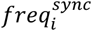 is the spiking frequency when stimulated with synchronized input and *freq*^*sparse*^ is the spiking frequency when stimulated with sparse input. *i* is the corresponding relevant current (*I*KA, *I*_CaT_ or *I*_NaT_).

Overall, the results from Fig. 9 show a relative frequency below 40%, suggesting that under the investigated conditions, the dendrite mainly behaves as an integrator.

## Discussion

This study examines an important aspect of the neural coding by focusing on different modes of *in vivo* dendritic synaptic integration. We show the role of inactivating currents on dendritic temporal properties, and how the modes vary from integration of synaptic potentials over time (long τ_m_) to coincidence detection of closely timed potentials (short τ_m_) (Mathews et al. 2010; Prescott and De Koninck 2005; Ceballos et al. 2018; Farries et al. 2010; Branco et al. 2016).

Our results point to the role of A-type potassium currents in shortening τ_m_, while T-type calcium and transient sodium currents lengthen it, similar to non-inactivating currents. This suggests inactivation reduces current strength without altering its profile. Additionally, activation and inactivation affect τ_m_ in opposite ways, with currents that shorten τ_m_ upon activation prolonging it upon inactivation and vice versa, consistent with previous studies (Richardson et al. 2003). These findings also apply to fast-activating, non-inactivating currents, e.g. M-type potassium and persistent sodium currents.

In terms of firing, we found that A-type potassium currents consistently reduce firing rates, whereas T-type calcium and transient sodium currents increase them. This is due to potassium currents hyperpolarizing the membrane, reducing voltage amplitude, and shortening temporal changes, which decreases excitability. In contrast, calcium and sodium currents depolarize the membrane, enhance voltage amplitude, and extend temporal changes, thereby increasing excitability.

In relation to the effects of inactivating currents under *in vivo*-like conditions with oscillatory inputs and synaptic fluctuations, both *I*_KA_ and *I*_NaT_, were observed scaling changes above -60 mV, showing monotonic trends with variability. The rise and decay phases followed distinct pathways, indicating that these currents differentiate between depolarization and hyperpolarization. No clear trend was found for *I*_CaT_, but its amplification pattern remained.

Activation and inactivation trajectories were linearly correlated for *I*_KA_ and *I*_CaT_, but nonlinear for *I*_NaT_, independent of kinetics. Surprisingly, activation/inactivation kinetics had minimal impact on voltage changes, with differences of less than 1 mV, suggesting potential misinterpretations when extrapolating *in vitro* data to *in vivo* contexts. This underscores the importance of *in vivo* recordings to fully understand dendritic behavior (Moore et al. 2017; Ujfalussy et al. 2018; Dembrow et al. 2022; Liao et al. 2024).

We can regard our results as a step towards understanding how dendrites process information. In particular, we have also tested temporally distinct patterns. Synaptic summation in the presence of A-type potassium currents promoted firing with synchronized input, while T-type calcium and transient sodium currents supported firing with sparse input. Thus, A-type potassium currents enhance synchrony-coding, whereas calcium and sodium currents favor rate-coding. These currents modulate dendritic integration in cortical and hippocampal dendrites (Stuart et al. 2015). Our findings align with experimental evidence showing that inactivating T-type calcium currents extend synaptic potentials (Connelly et al. 2016; Gillessen and Alzheimer 1997; Liu and Shipley 2008; Prescott and De Koninck 2005; Urban et al. 1998), while inactivating A-type potassium channels shorten them (Hoffman et al. 1997; Mathews et al. 2010). Notably, we are the first to show that transient sodium currents have a lengthening effect.

Our results align with previous computational simulations. For instance, Remme et al. (2011) found that EPSP amplitude and duration decrease due to low-voltage-activated potassium currents and increase due to persistent sodium currents. Ceballos et al. (2017b) reported that persistent sodium current amplifies and prolongs EPSPs in a voltage-dependent manner. Hoffman et al. (1997) showed that A-type potassium current activation reduces EPSP amplitude in CA1 pyramidal neuron dendrites. Branco et al. (2016) demonstrated that balanced persistent sodium and potassium currents can generate long-lasting EPSPs, enabling near-perfect synaptic integration. Additionally, Carter et al. (2012) used computational models to predict that sodium currents mediate subthreshold transient current activation by EPSPs. The inactivating A-type potassium and T-type calcium currents have also been studied in simulations in the context of resonance (Rathour and Narayanan, 2012). However, to our knowledge, we are the first to investigate their role in dendritic spike integration modes under *in-vivo*-like conditions.

Future work should explore other factors influencing the activation/inactivation properties of ionic currents and systematize findings across species. For example, sodium and potassium current voltage-dependence is more depolarized in humans than mice (Wilbers 2023), and isoflurane hyperpolarizes sodium current inactivation (Qiu et al. 2023). Neuromodulation also enhances sodium current inactivation (Chen et al. 2006). Though we focus on *in vivo*-like states, future studies could investigate how alternating states affect synaptic integration.

## Author contributions

C.C.C. and R.F.O.P. designed the research. C.C.C. carried out all simulations and analyzed the data. C.C.C. and R.F.O.P. wrote the article.

