## Supporting Material for "Dendritic synaptic integration modes under *in vivo*-like states"

**Supplementary Material - Dendritic synaptic integration modes under *in vivo*-like states**
Cesar C. Ceballos^1^, Rodrigo F. O. Pena^1,2 *
1^ Department of Biological Sciences, Florida Atlantic University, Jupiter, FL 33458, USA
^2^ Stiles-Nicholson Brain Institute, Florida Atlantic University, Jupiter, FL 33458, USA
***** Corresponding author
E- mail:

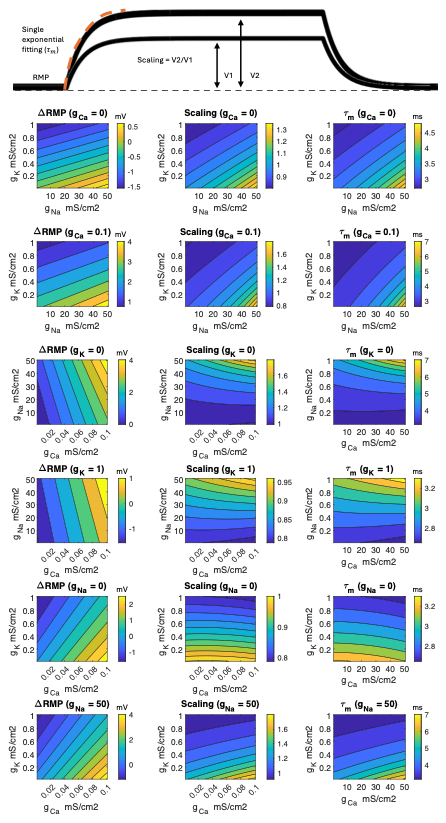

Fig S1. Heatmaps of difference of resting membrane potentials (ΔRMP), scaling and membrane time constant (τ_m_). Dendritic model contains all 3 currents (I_KA_, I_CaT_ and I_NaT_) plus leak current. In each heatmap two conductance values were varied and one was fixed. ΔRMP was calculated as the subtraction between RMPs of both traces. Scaling was calculated as the ratio of the voltages V_2_/V_1_. Membrane time constant was obtained from a fitting of the rise phase using a single exponential function.

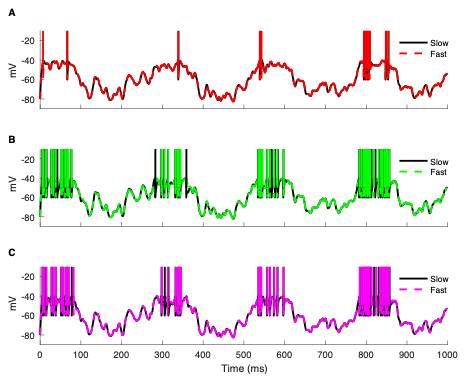

Fig S2. Comparison of spiking activity using actual kinetics vs fast kinetics (i.e. τ_act_ = τ_inact_ = 0.1 ms). A. *I*_KA_. B. *I*_CaT_. C. *I*_NaT_. The conductance values were: g_KA_ = 1, g_CaT_ = 0.1 and g_NaT_ = 50, respectively.

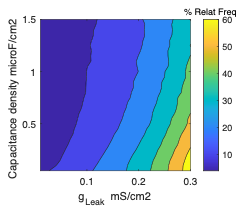

Fig S3. Heatmaps of % relative frequency (% relat freq) computed as difference between the sparse vs synchronized synaptic inputs spanned over different leak conductance values (x axis) vs capacitance density (y axis). The % relative frequency is an indicator of whether the dendrite acts as an integrator of coincidence detector (100% for perfect coincident detector and 0% for perfect integrator). $\% relative frequency=100\frac{({freq}^{sync}-{freq}^{sparse})}{{freq}^{sparse}}$, where ${freq}^{sync}$ is the spiking frequency when stimulated with synchronized input and ${freq}^{sparse}$ is the spiking frequency when stimulated with sparse input.
